# Gene flow influences the genomic architecture of local adaptation in six riverine fish species

**DOI:** 10.1101/2021.05.18.444736

**Authors:** Yue Shi, Kristen L. Bouska, Garrett J. McKinney, William Dokai, Andrew Bartels, Megan V. McPhee, Wesley A. Larson

## Abstract

Understanding how gene flow influences adaptive divergence is important for predicting adaptive responses. Theoretical studies suggest that when gene flow is high, clustering of adaptive genes in fewer genomic regions would protect adaptive alleles from among-population recombination and thus be selected for, but few studies have tested this hypothesis with empirical data. Here, we used RADseq to generate genomic data for six fish species with contrasting life histories from six reaches of the Upper Mississippi River System, USA. We then conducted genome scans for genomic islands of divergence to examine the distribution of adaptive loci and investigated whether these loci were found in inversions. We found that gene flow varied among species, and adaptive loci were clustered more tightly in species with higher gene flow. For example, the two species with the highest overall *F*_ST_ (0.03 - 0.07) and therefore lowest gene flow showed little evidence of clusters of adaptive loci, with adaptive loci spread uniformly across the genome. In contrast, nearly all adaptive loci in the species with the lowest *F*_ST_ (0.0004) were found in a single large putative inversion. Two other species with intermediate gene flow (*F*_ST_ ~ 0.004) also showed clustered genomic architectures, with most islands of divergence clustered on a few chromosomes. These results provide important empirical evidence to support the hypothesis that increasingly clustered architectures of local adaptation are associated with high gene flow. Our study utilized a unique system with species spanning a large gradient of life histories to highlight the importance of gene flow in shaping adaptive divergence.

## Introduction

Understanding the genomic basis of adaptation is a central goal of evolutionary biology. Research on this topic largely focuses on identifying genetic markers involved in adaptation and assessing the distribution of these markers across the genome (Narum & Hess 2011; Yeaman 2013; Lotterhos & Whitlock 2014; Hoban *et al*. 2016; Forester *et al*. 2018). Substantial efforts have focused on this area of research for decades (Smith & Haigh 1974; Rieseberg 2001; Noor *et al*. 2001). However, results have been highly variable across taxa and systems, making it difficult to gain a mechanistic understanding of the evolutionary processes that influence the genomic landscape of adaptation. For example, many studies have found that alleles contributing to local adaptation tend to be clustered together in genomic islands of differentiation, while other studies have found little or no evidence of adaptive alleles clustering within genomic islands (Nosil *et al*. 2009; Strasburg *et al*. 2012; Roda *et al*. 2017; Johannesson *et al*. 2020; Thompson *et al*. 2020). This mixed evidence raises an important evolutionary question: when are loci affecting adaptive divergence expected to be tightly clustered?

Interpreting results from genome scans in the context of gene flow may aid in the understanding of genomic landscapes of adaptation (Marques *et al*. 2016). Gene flow can be beneficial for maintaining population connectivity and genetic diversity by introducing novel genetic variation but it can also impede local adaptation by introducing maladaptive foreign alleles into a locally adapted populations (Bolnick & Nosil 2007). One potential evolutionary ‘solution’ that may minimize maladaptive effects of gene flow is for selection to favor clustered architectures of adaptation, where adaptive alleles are tightly linked and locally favorable combinations of alleles are protected from disruption via low recombination (Yeaman 2013; Roesti 2018).

Several mechanisms have been proposed to explain the observations of clustered genomic architectures of adaptive alleles when gene flow is high, including divergence hitchhiking and the utilization of genomic rearrangements to protect adaptive loci from recombination. Divergence hitchhiking occurs when gene exchange between diverging populations is reduced around a gene under strong divergent selection (Via 2012). This process can produce islands of differentiation spanning multiple megabases, as free recombination among populations is reduced due to assortative mating (Via 2012). Genomic rearrangements, such as chromosomal inversions, can also facilitate adaptation in the face of high gene flow and lead to genomic islands of differentiation (Hoffmann & Rieseberg 2008; Yeaman 2013; Tigano & Friesen 2016; Roesti 2018; Wellenreuther & Bernatchez 2018; Aguirre Liguori *et al*. 2019; Huang *et al*. 2020; Cayuela *et al*. 2020). Inversions are generally not deleterious and do not impact gene function unless the inversion breakpoint occurs within a gene (Faria *et al*. 2019). However, recombination between inverted and noninverted arrangements is rare as recombinant gametes are generally inviable (Huang & Rieseberg 2020). Therefore, if an inversion isolates multiple adaptive alleles, this architecture will likely be favored, because co-adapted genotypes will be protected from recombination and allowed to evolve independently even in high gene flow environments (Rogers *et al*. 2013; Yeaman 2013).

Although the theories described above posit that the rate of evolution towards clustered architectures of local adaptation should increase with gene flow, this hypothesis has largely been tested with simulations rather than empirical data. For example, Yeaman & Whitlock (2011) used simulations to demonstrate increasing migration rate, or *m*, leads to increasingly concentrated genomic architectures of adaptation. However, when *m* is too high, adaptive divergence is unlikely because frequent migration prevents even a perfectly adapted mutation from overcoming the homogenizing effects of gene flow. A subsequent simulation study (Yeaman 2013) highlighted that genomic rearrangement may often be an important component of local adaptation and when genomic rearrangements are present, tight clustering of adaptive loci can readily evolve even with high *m*.

In this study, we investigate how gene flow influences the genomic architecture of adaptation using genomic data from six riverine fish species that encompass a diverse suite of life histories and dispersal potentials (Figure 1B). These fish were sampled from the same sites in the Upper Mississippi River System (UMRS) in the midwestern United States. The UMRS is an interconnected large river system that hosts a diversity of aquatic habitats in terms of temperature, turbidity, productivity and flow (Figure 1A & C). Our study system provides a unique opportunity to compare the genomic architecture of local adaptation in a natural environment for species with contrasting life histories and assess the influence of gene flow on genomic architecture. Specifically, we test the hypothesis that the genomic islands of differentiation are less frequent but larger for species with relatively high gene flow, whereas genomic islands are more numerous and dispersed throughout the genome for species with low degrees of gene flow. Our multi-species approach investigating six species inhabiting the same environments is unique, as most previous studies have focused on closely related species pairs or ecotypes (Nadeau *et al*. 2012; Renaut *et al*. 2012) rather than divergent species inhabiting the same environments.

**Figure 1.**
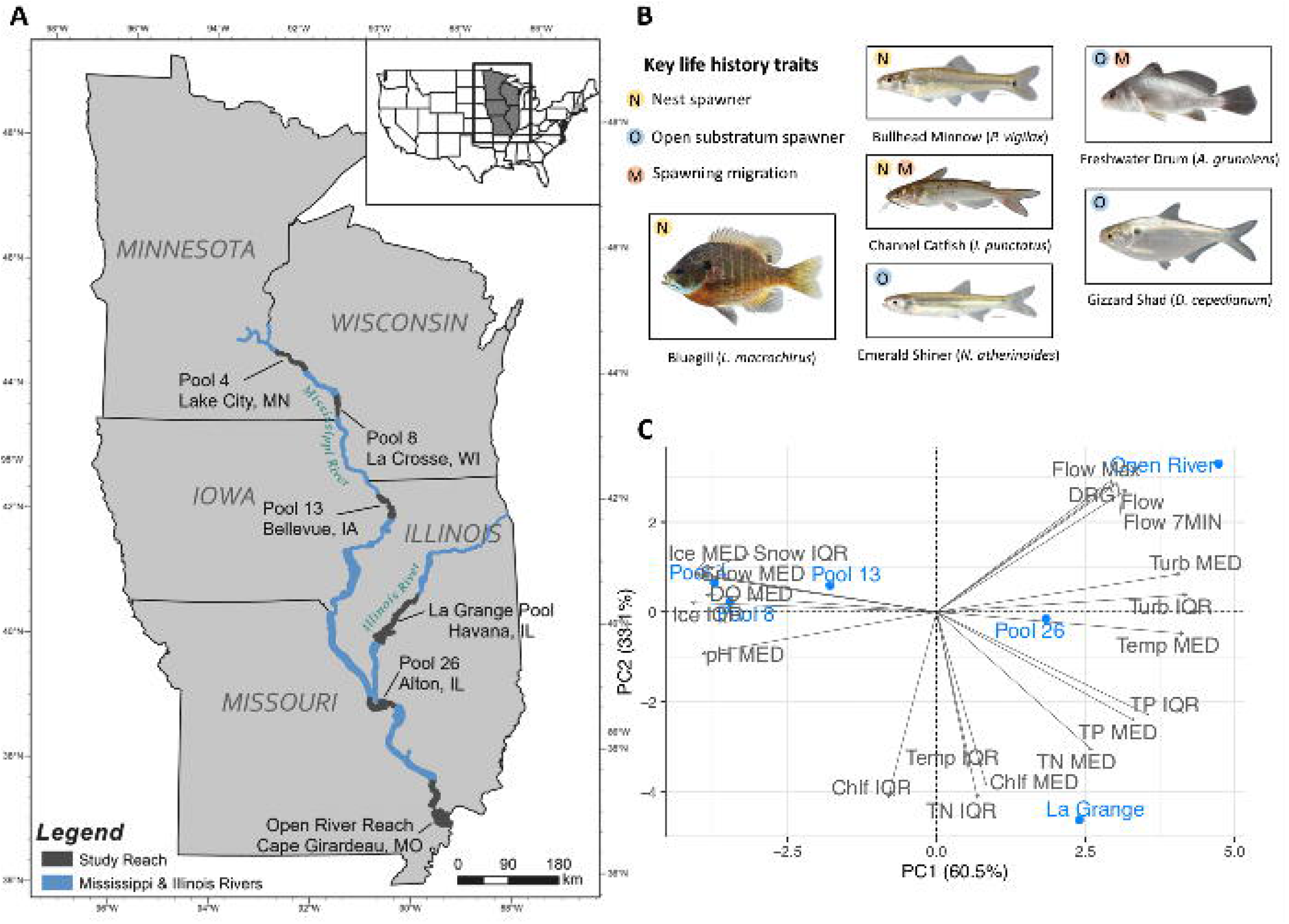
(A) Map of the six study reaches along the Upper Mississippi River System, (B) key reproduction-related life history traits of the six study species, and (C) positions of the six study reaches in the environmental space of 20 variables using PCA biplot. See Table S1 for details of life history traits and Table S2 for details of environmental data. Use of fish images is permitted by Uland Thomas.

## Materials and Methods

### Study Design and Genotyping

We collected genetic samples from six fish species found in the UMRS which are native to and commonly found in the region and have not been extensively stocked: Bullhead Minnow (*Pimephales vigilax*), Bluegill (*Lepomis macrochirus*), Freshwater Drum (*Aplodinotus grunniens*), Channel Catfish (*Ictalurus punctatus*), Gizzard Shad (*Dorosoma cepedianum*), and Emerald Shiner (*Notropis atherinoides*). The UMRS is congressionally defined as the commercially navigable portions of the Mississippi River main stem north of Cairo, Illinois” and commercially navigable tributaries, including the entire Illinois River (Water Resources Development Act of 1986, 33 U.S.C. §§ 652). Fin-clip samples were collected from adult fish in summer 2018 and 2019 across six river reaches (Figure 1A); five of the study reaches are navigation pools, named for their downstream lock and dam, and the other study reach (Open River Reach) is an unobstructed, channelized reach. We targeted a sample size of at least 48 samples per species per reach. Samples were genotyped at thousands of SNPs using restriction site-associated DNA (RAD) sequencing (see Supplementary Methods). Data on life history traits for each species, including exploitation status, feeding guild, habitat guild, reproductive guild, spawning migration, and total length were summarized in Table S1. We also obtained data for 20 environmental variables across the six river reaches (Table S2).

### Identification of GEA Outliers and Putatively Neutral SNPs

Recent studies have suggested that genotype-environment association (GEA) methods are more robust for identifying adaptive loci than traditional differentiation-based methods (Rellstab *et al*. 2015; Forester *et al*. 2018). Differentiation-based outlier tests identify loci with high *F*_ST_ values, which are expected for loci involved in hard selective sweeps with large changes in allele frequencies (Brauer *et al*. 2016; Forester *et al*. 2018). By comparison, GEA analyses identify genetic variants associated with particular environmental factors and can identify loci under polygenic and “soft” selective sweeps with relatively small changes in allele frequencies, providing a more complete view of the genomic landscape of adaptation (Eckert *et al*. 2010; Brauer *et al*. 2016; Forester *et al*. 2018). For these reasons, we focused on identifying putatively adaptive loci (henceforth “adaptive loci”) using three GEA methods: redundancy analysis, latent factor mixed models, and a Bayesian method (Bayenv2). Details of these methods can be found in the Supplementary Methods. Prior to all three GEA analyses, we conducted principal component analysis (PCA) on 20 standardized environmental variables. Based on Kaiser-Guttmann criterion and the broken stick model, we retained the first two significant PCs as environmental composite variables in order to remove collinearity among variables (Figure S1A). Variables related to temperature, turbidity, pH, and dissolved oxygen had high loadings on environmental PC1 (Figure S1B), whereas productivity and flow-related variables contributed significantly to environmental PC2 (Figure S1C). We defined putatively adaptive SNPs as the GEA outliers (henceforth “GEA outliers”) identified by at least two GEA methods. To determine which environmental PC that each GEA outlier was most strongly correlated with, we compared correlation coefficients between each environmental PC and genotype for each outlier using R function *cor* and assessed which environmental PC had the highest correlation coefficient.

To identify datasets of putatively neutral SNPs (henceforth “neutral SNPs”) for each species, we combined results from GEA analyses with results from additional differentiation-based outlier tests. While differentiation-based outlier tests may produce a large number of false positives, they are still useful for conservatively identifying neutral SNPs (Holderegger *et al*. 2006). Therefore, we ran Bayescan, Arlequin, OutFLANK and *pcadapt* on each species (see Supplementary Methods for details). We defined neutral SNPs as those that were not identified as outliers by any of the aforementioned seven methods.

### Neutral Genetic Differentiation

We used three methods to estimate population structure for each species with their neutral datasets. First, we calculated global Fstp (*F*_ST_ corrected for sampling bias) using the function *basic.stats* in *hierfstat* v.0.04-22 (Goudet 2005). Next, we calculated *F*_ST_ between all pairs of river reaches using *genet.dist* function (method=”WC84”) in *hierfstat*. Significance was assessed by calculating 95% confidence interval of pairwise *F*_ST_ values using *boot.ppfst* function (nboot=1000) in *hierfstat*. A pairwise *F*_ST_ value was considered significant if its confidence interval did not include zero. Lastly, we conducted PCA implemented in the R package *adegenet* v2.1.2 (Jombart 2008) to investigate genetic differentiation among individuals.

### Genome Scans for Genomic Islands of Differentiation

We aligned SNPs to reference genomes and conducted genome scans to investigate the genomic landscape of adaptive divergence. Channel Catfish is the only species with a high-quality reference genome available in our study. For the other five species, we used high-quality reference genomes (chromosome-level assemblies) from closely related species (Table S3). Sequences of filtered RAD loci were mapped to reference genomes with BWA-MEM v 0.7.17 using default settings (Li 2013). We retained sequences with mapping quality > 20 and removed sequences with “SA:Z” (chimeric alignment) and “XA:Z” tags (alternative hits) using *SAMtools* v1.10 (Li *et al*. 2009).

To identify genomic islands of differentiation, we first calculated Fstp per locus using the *basic.stats* function in *hierfstat* for all aligned SNPs across genomes. We then used a Hidden Markov Model (HMM) approach implemented in the R package *HiddenMarkov* v.1.8-11 (Hofer *et al*. 2012) to assign each SNP to one of three underlying states, “genomic background”, “regions of high differentiation” and “regions of low differentiation” based on their Fstp values, following the methods detailed in Marques *et al*. (2016). Each of these identified regions can consist of one or many consecutive SNPs depending on the landscape of differentiation.

The HMM approach identified a large number of highly differentiated regions (i.e., genomic islands of differentiation), but many did not show especially high levels of differentiation and may be false positives. Therefore, we chose to retain only the genomic islands that contained at least one differentiation outlier SNP identified by Bayescan, Arlequin or OutFLANK. We excluded outliers identified only by *pcaadpt* because we discovered this method identified a much higher number of outliers compared to other methods, which could potentially increase false positive rate for island detection. We removed genomic islands in situations where a chromosome only had one island and this island had only one SNP. We also removed islands located on unplaced scaffolds.

### Identification and Analysis of Putative Inversions

To identify putative inversions, we conducted a sliding window analysis of population structure across genomes using the R package *lostruct* (Li & Ralph 2019) following the methods described in Huang *et al*. (2020). We replaced missing genotypes with the most frequent genotype and divided each genome into nonoverlapping windows of either 20 or 50 SNPs depending on the total number of aligned SNPs for each species. We then used a 40-dimension space multidimensional scaling (MDS) analysis to measure the differences in population structure patterns among windows, and we defined outlier windows as those with absolute values of loadings greater than 4 standard deviations above the mean averaged across all windows in the genome (Huang *et al*. 2020). Outlier windows (single or consecutive) were candidate regions for putative inversions. We also conducted three additional analyses on putative inversion regions to provide additional evidence of inversions as suggested by Huang *et al*. (2020). First, because inversions only suppress recombination in heterozygotes, three distinct genotypic clusters (0, 1, 2) should be detected along PC1 using PCA, with the outside clusters (0 and 2) representing two homozygous groups for alternative orientations and the middle cluster (1) representing the heterozygous group between inversion haplotypes (McKinney *et al*. 2020). The discreteness of the clustering was calculated as the proportion of the between-cluster sum of squares over the total using the R function *kmeans* in *adegenet*. Second, we compared heterozygosity (the proportion of heterozygotes) among three clusters identified by PCA using Wilcoxon tests (α = 0.05) to further confirm the middle group had significantly higher heterozygosity. Finally, we calculated linkage disequilibrium, or LD (r^2^) using PLINK v1.9 (Purcell *et al*. 2007; Chang *et al*. 2015) for SNPs with MAF > 0.01 on chromosomes with outlier windows and compared r^2^ with all samples to r^2^ calculated only from samples that were homozygous for the most common orientation. Since inversions are expected to only suppress recombination in heterokaryotypes, recombination in homokaryotypes should be unaffected.

We considered a region as a putative inversion only if all of the following criteria were met: (1) a distinct three-cluster PCA pattern with discreteness > 0.9; (2) significantly elevated heterozygosity in the middle PCA cluster compared to the other two clusters; and (3) elevated LD calculated with all samples, but not with homozygous samples. We assumed that the more derived inversion arrangement would have lower heterozygosity given its relatively recent origin compared to the ancestral state (Laayouni 2003; Twyford & Friedman 2015; Knief *et al*. 2016). Notably, when examining our data, we found five additional regions with discreteness very close to 0.9 (0.893 - 0.898) that displayed distinct three-cluster PCR patterns, and we included these regions as candidates for putative inversions as well.

To investigate patterns of population structure at putative inversions, we calculated genotype frequencies in each river reach for each putative inversion. Additionally, we conducted PCA analyses using all SNPs that were successfully aligned to genomes, SNPs within the identified inversions, and the remaining aligned SNPs after the SNPs in putative inversions were removed to compare patterns of genetic structure inferred from datasets including and not including putative inversions.

### Identification and Analyses of Large Clusters of Adaptive Loci

Adaptive loci can be found across many areas of the genome or can be concentrated (i.e., clustered) in only a few genomic regions. To determine whether the genomes of our species contained clustered architectures of adaptative loci, we investigated the distribution of GEA outliers and islands of differentiation identified by HMM across the genome. We defined chromosomes exhibiting clustered architecture of adaptation as chromosomes containing at least 3 GEA outliers or 20% of the total HMM islands within a given species. We then calculated the following genetic metrics to characterize the genomic properties of clustered architectures that we observed: Fstp, heterozygosity (*H_O_*), absolute differentiation (*D_xy_*), and linkage disequilibrium (LD).

Fstp and *H_O_* were calculated using the *basic.stats* function in *hierfstat* as described previously. Pairwise per-site *D_xy_* was calculated as *p*_1_(1 ‒ *p*_2_) + *p*_2_(1 – *p*_1_), where *p*_1_ is the frequency of a given allele in the first population and *p*_2_ is the frequency of that allele in the second population (Irwin *et al*. 2016). Allele frequency was estimated using *makefreq* function (missing = “mean”) in *adegenet*. Overall *D_xy_* was calculated as the mean of all pairwise *D_xy_* values. LD (r^2^) was calculated using PLINK v1.9 for SNPs with MAF > 0.01. We included *D_xy_*, an absolute measure of genetic differentiation, because defining adaptive genomic regions based solely on relative measures of differentiation, such as Fstp, may identify regions resulting from processes other than adaptation (Cruickshank & Hahn 2014).

For each chromosome, we used Wilcoxon tests (α = 0.05) to test for significant differences in genetic metrics between SNPs within the islands and SNPs outside the islands (chromosomal background). We also visualized differences in these metrics with boxplots. To ensure the differences we observed in four genetic metrics were not due to island size, we randomly selected five windows outside of the islands as chromosomal background with window size (number of SNPs) set as the average size of all HMM islands found on the corresponding chromosomes. When calculating the average size of the HMM islands, we removed the HMM islands containing only one SNP to avoid downward biasing the window size of random windows in the chromosomal background. We also made sure the randomly selected windows spanned similar distance (± 10% bp) compared to the average of all HMM islands found on the same chromosomes.

Lastly, we conducted Gene Ontology (GO) enrichment tests to test for functional enrichment of genes in the HMM islands located within the six chromosomes displaying clustered architecture. We extracted genes within 10 Kb of a SNP for all SNPs located within the islands for all six chromosomes, except for chromosome 9 in Emerald Shiner. Since all of the HMM islands on chromosome 9 in Emerald Shiner were clustered inside of the identified inversion and there was a relatively smaller number of aligned loci, we extracted genes within 20 Kb of a SNP for all SNPs located within the identified inversion on the chromosome 9 in Emerald Shiner instead. See Supplementary Materials for detailed methods about GO enrichment tests.

## Results

### Summary of Sequencing, GEA Outliers, and Neutral SNPs

We RAD sequenced a total of 1,712 individuals, ranging from 275 - 288 individuals per species. RAD sequencing yielded an average of 5,780,907 retained reads per individual (range = 16,799 - 47,250,859). After filtering, 1,417 individuals (179 - 256 individuals per species) were retained and genotyped at 10,834 - 28,313 polymorphic SNPs depending on the species (Table S3). Out of these polymorphic SNPs, 0.04 % to 0.3% were identified as GEA outliers, and 95.9 % - 99.4% were identified as neutral SNPs in each species (Table S4). For most species, the majority of GEA outliers were found to be strongly associated with environmental PC1 (temperature, turbidity, pH, and dissolved oxygen related). In contrast, GEA outliers in Freshwater Drum were strongly associated with environmental PC2 (productivity and flow related) (Table S4).

### Neutral Genetic Differentiation

Patterns of population structure estimated from the neutral datasets spanned a large gradient of genetic differentiation across species (Figure 2, Table S5). Bullhead Minnow had the highest global Fstp value of 0.0720 with pairwise *F*_ST_ values ranging from 0.0041 to 0.1543, followed by Bluegill (global Fstp = 0.0302; pairwise *F*_ST_ = 0.0014 - 0.0739), Freshwater Drum (global Fstp = 0.0050, pairwise *F*_ST_ = −0.0003 - 0.0169), Channel Catfish (global Fstp = 0.0025, pairwise *F*_ST_= 0.0003 - 0.0048), and Gizzard Shad (global Fstp = 0.0024, pairwise *F*_ST_ =0.0003 - 0.0051). Emerald Shiner had the lowest global Fstp value among all six species, 0.0004, with pairwise *F*_ST_ values ranging from −0.0003 to 0.0016.

**Figure 2.**
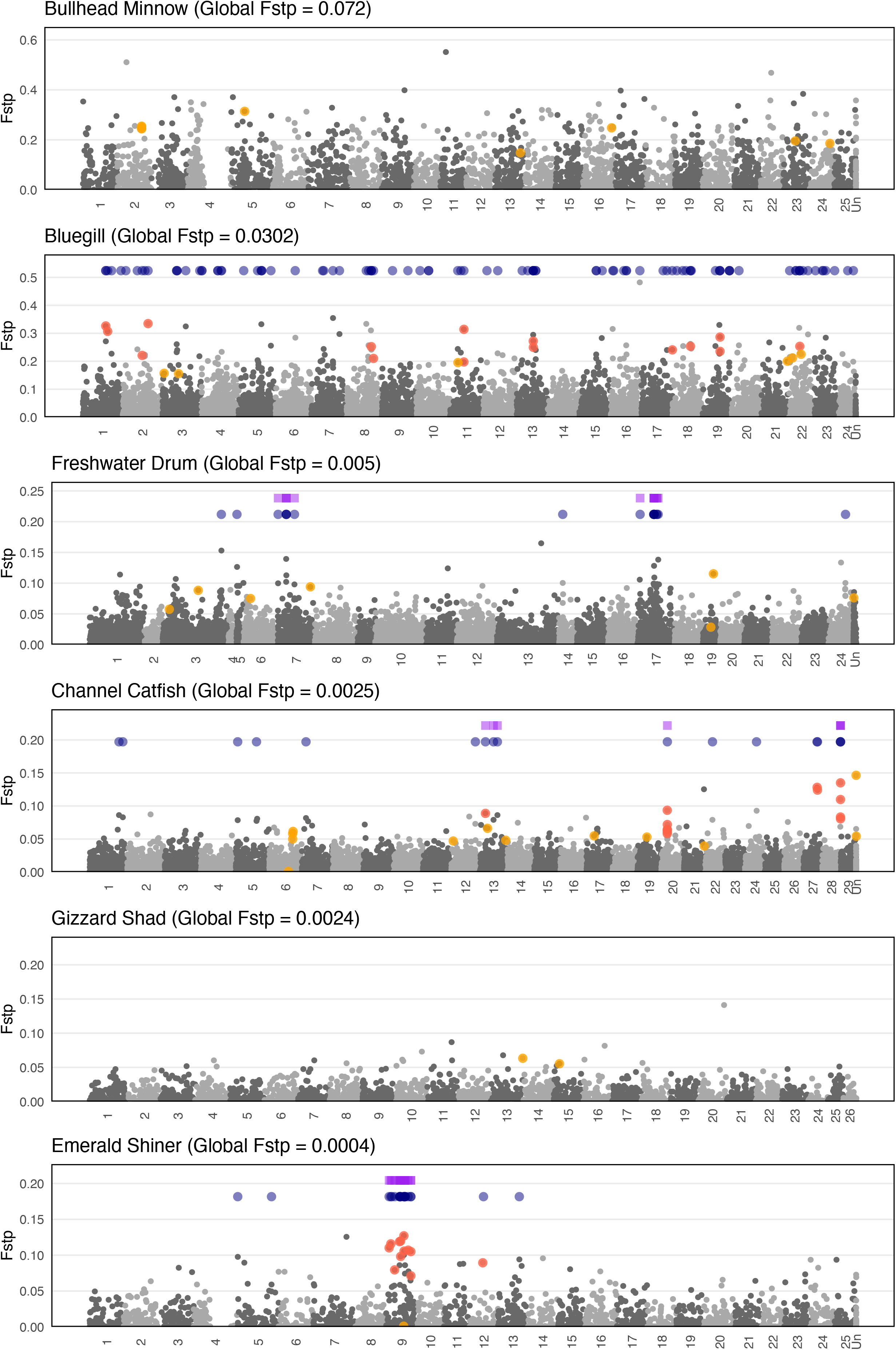
Manhattan plots depicting the genomic landscape of differentiation (Fstp, corrected *F_ST_*) across the genomes for the six study species. Species are ordered based on neutral population differentiation, with neutral global Fstp values labeled next to the species name. At the top of each plot, genomic islands of differentiation identified using HMM after filtering are in blue, and islands located within the chromosomes showing clustered architecture are in purple. GEA outliers found within islands are in red, whereas those found outside of islands of genomic divergence are in orange. Reference genomes and alignment summary can be found in Table S3.

Results of the PCAs (Figure 3) corroborated the patterns described above. In Bullhead Minnow, we detected five genetic clusters, with individuals from each river reach forming a single cluster except for Pool 8 and Pool 13, which were grouped together. In Bluegill, individuals from the three northern river reaches (Pool 4, Pool 8, and Pool 13) were genetically similar, Pool 26 and La Grange formed a second cluster, while the most southerly reach, Open River, formed its own cluster. In Freshwater Drum, individuals from La Grange grouped separately from other populations along with some individuals from Pool 26 and Open River. In Channel Catfish, individuals from the Open River were slightly separated from all the other reaches. Lastly, Gizzard Shad and Emerald Shiner showed no apparent population structure.

**Figure 3.**
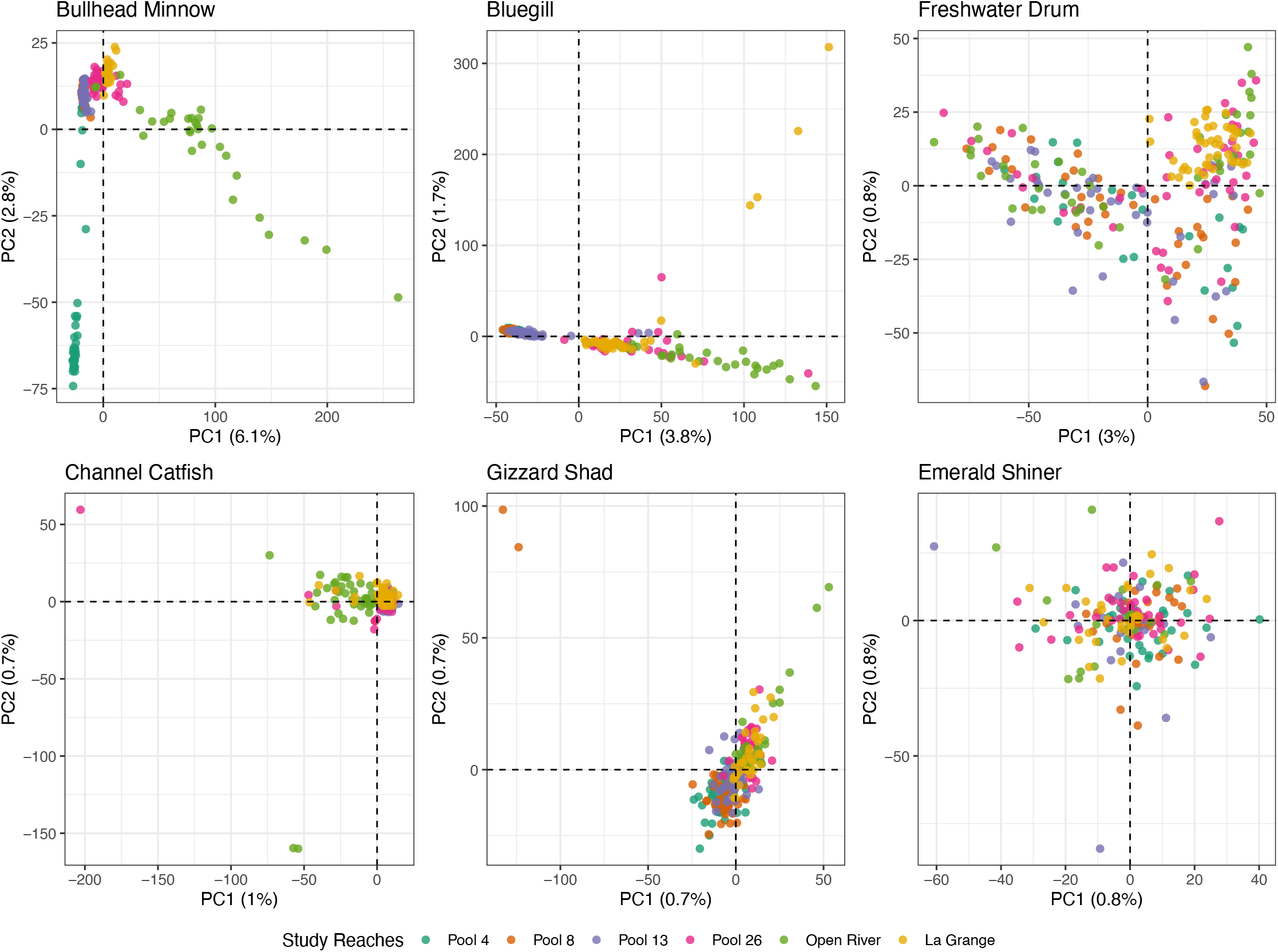
Principal component analyses using neutral SNPs only for the six study species. The percentage of variance explained by each principal component (PC) is labeled on the *x*- and *y*- axes.

### Genome Scan for Genomic Islands of Differentiation

We aligned SNPs to reference genomes and conducted genome scans to investigate the genomic landscape of adaptive divergence. A total of 3,348 - 16,620 loci were aligned to the corresponding reference genomes with alignment rate varying from 26.4% to 97.5% depending on genetic divergence from the reference species (Table S3). Correspondingly, a total of 2 - 25 GEA outliers were aligned with alignment rate per species varying from 25.7% to 100% (Table S4).

Genome scan results revealed highly variable genomic landscapes of population differentiation among the six species (Figure 2). In general, GEA outliers and HMM islands in species with lower neutral differentiation were more tightly clustered and found on fewer chromosomes, whereas those in species with higher neutral differentiation were spread out across the genome. Bullhead Minnow (highest neutral population structure) displayed a high level of baseline differentiation without obvious peaks of highly differentiated loci. We only detected 2 islands on 2 chromosomes and there were no GEA outliers located within the islands. In Bluegill, the species with the second highest neutral population structure, we identified 83 islands that were dispersed across nearly all chromosomes (22/24) with no chromosomes containing more than 8% of islands. Additionally, 15 out of 21 aligned GEA outliers (71%) were located in 12 islands across 9 chromosomes with only 3 islands having more than one GEA outliers (up to 2). Freshwater Drum had an intermediate level of population differentiation and displayed a more clustered architecture of genomics islands of differentiation compared to Bullhead Minnow and Bluegill. In total, 14 islands were detected across 6 chromosomes and no GEA outliers were found within islands. Of these islands, 3 islands (21%) were clustered on chromosome 7 and 7 islands (50%) were clustered on chromosome 17. Channel Catfish had a relatively low level of differentiation and displayed highly clustered architectures of adaptation. We identified 15 islands across 10 chromosomes with 3 islands (20%) clustered on chromosome 13. Out of 25 aligned GEA outliers, 6 (24%) were located on an island on chromosome 20 and 4 (16%) were located on an island on chromosome 28. Gizzard Shad had a similar level of neutral global Fstp as Channel Catfish, but we did not detect any islands of high differentiation, possibly due to its relatively low genome alignment rate (26.4%). Lastly, Emerald Shiner, the species with lowest overall neutral population differentiation, displayed the strongest signal of clustered architecture of local adaptation. In Emerald Shiner, 15 islands were detected across 4 chromosomes, of which, 11 (73%) were clustered on chromosome 9. A total of 11 out of 13 aligned GEA outliers (85%) were found on chromosome 9.

### Identification and Analyses of Putative Inversions

Using local PCA in *lostruct*, we identified 21 candidate regions for putative inversions where individuals clustered into three distinct groups on PC1 and with the middle PCA cluster displaying significantly higher heterozygosity than the other two clusters (Table S6). Of all candidate regions, only the ones on chromosome 14 in Channel Catfish and chromosome 6, 9, and 19 in Emerald Shiner were characterized by elevated LD blocks extending over several Mb, while LD decayed very quickly on other chromosomes (Figure S2). However, we detected recombination suppression in both heterozygous and homozygous groups in the outlier region on chromosome 14 in Channel Catfish (Figure S3). This pattern is inconsistent with the theory that inversions should only suppress recombination in heterokaryotypes, so this region was excluded. Only the candidate regions on chromosome 6 (Figure S4), 9 (Figure 4), and 19 (Figure S5) in Emerald Shiner passed our stringent criteria and were considered as putative inversions. These three putative inversions spanned large genomic regions, 18.0, 42.7, and 25.6 Mbp, respectively (Table S6). The two homokaryotypes presented significant differences in heterozygosity for all three putative inversions (Figure 4B, S5B and S6B) and we assumed that the arrangement with lower heterozygosity was the derived inverted type. The putative inversion on chromosome 9 (cluster 0) was only detected in the three southern river reaches (Figure 4C), whereas the other two inversions occurred at similar frequency across all six river reaches, ranging in frequencies from 0.15 to 0.39 for the inversion on chromosome 6 (cluster 2; Figure S4C), and from 0.19 to 0.28 for the inversion on chromosome 19 (cluster 0; Figure S5C). Moreover, GEA outliers and HMM islands were consistently associated with the putative inversion on chromosome 9 in Emerald Shiner (Figure 2). However, no GEA outliers or HMM islands were found within the inversions on the chromosome 6 and 19.

**Figure 4.**
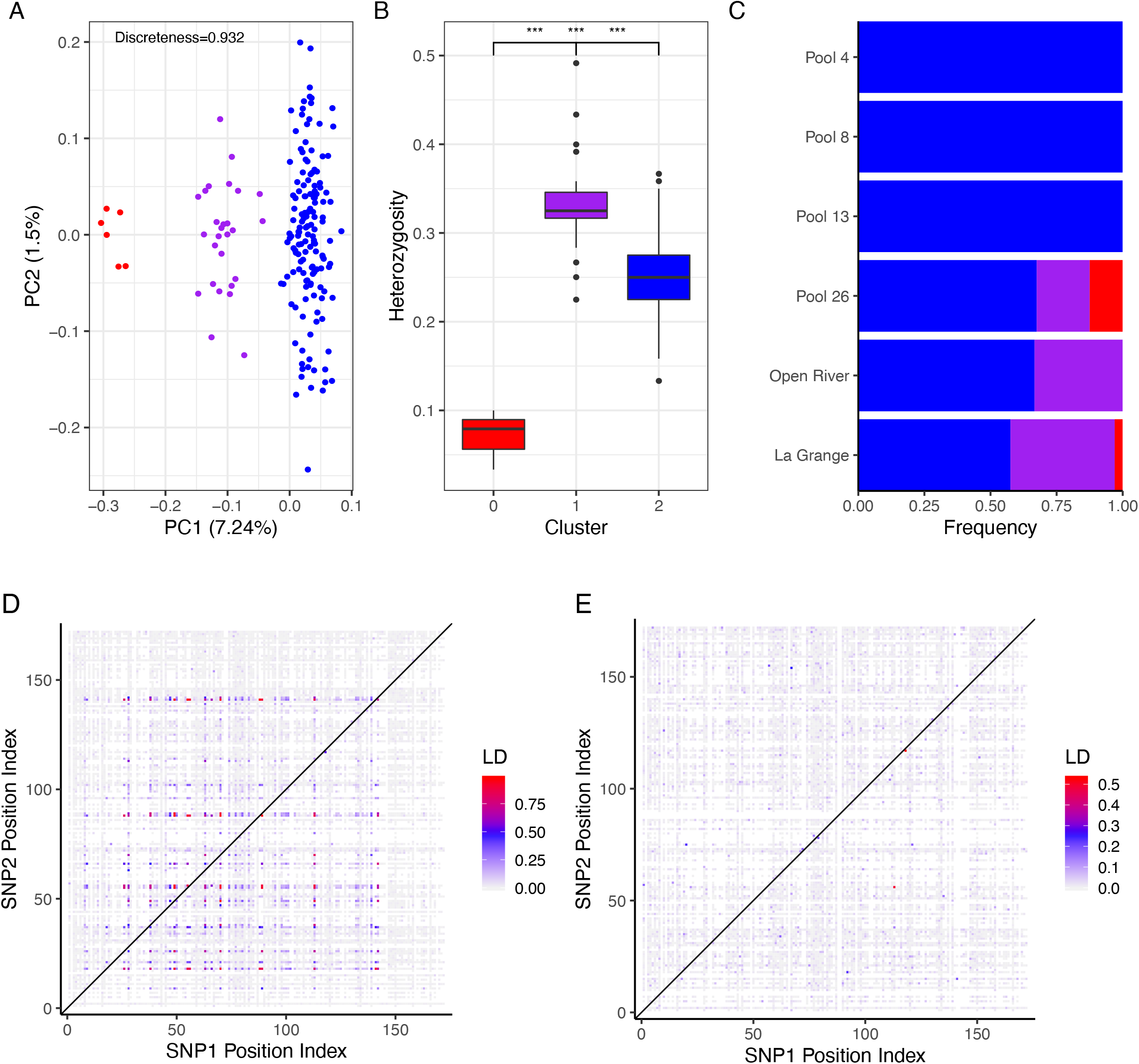
Characterization of putative inversion on chromosome 9 in Emerald Shiner. (A) PCA based on SNPs within the putative inversion region. Three clusters identified using k-means clustering correspond to two homozygote groups (blue and red) and a heterozygote group (purple). The discreteness of the clustering was calculated by the proportion of the between-cluster sum of squares over the total using the R function *kmeans* in *adegenet*. (B) Observed individual heterozygosity in each PCA cluster. Significance was assessed using Wilcoxon tests with alpha level of 0.05. Note: *** = 0.001. (C) Genotype frequency distribution for putative inversion across six study reaches. Bars represent the proportion of individuals belonging to a PCA cluster. (D) and (E) are LD heatmaps for chromosome 9 using all individuals (D) and only individuals homozygous for the more common orientation (E).

Analyzing datasets with and without putative inversions in Emerald Shiner produced substantially different patterns of genetic structure (Figure 5). Both PCA analyses based on all aligned SNPs and SNPs within the three identified inversions showed a similar genetic structure pattern, with six well-separated clusters. This illustrates that the clustering inferred from the full dataset is driven by these three inversions. After the SNPs in these inversions were removed, the remaining aligned loci demonstrated a lack of clustering, with panmictic population structure.

**Figure 5.**
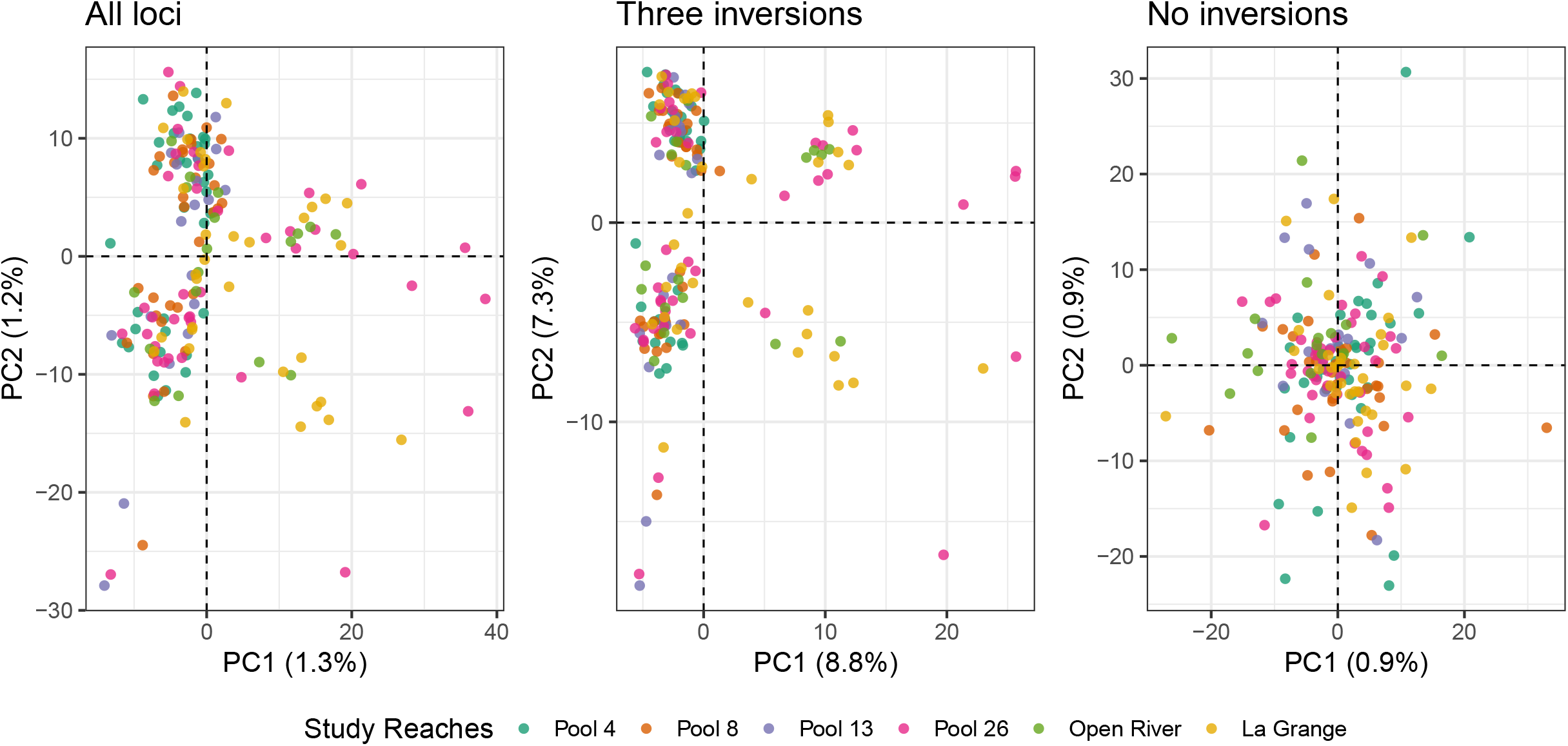
Principal component analyses for Emerald Shiner using different sets of loci: (A) All aligned SNPs (3,348 SNPs); (B) Putative inversions on chromosome 6, 9 and 19 (228 SNPs); (C) After the removal of three putative inversions (3,120 SNPs). The percentage of variance explained by each principal component (PC) is labeled on the *x*- and *y*- axes.

### Genomic Properties of Large Clusters of Adaptive Loci

The following chromosomes exhibited highly clustered architecture with at least 3 GEA outliers or 20% HMM islands within a given species: (1) chromosome 7 and 17 in Freshwater Drum; (2) chromosome 13, 20, and 28 in Channel Catfish; and (3) chromosome 9 in Emerald Shiner (Figure 2). The HMM islands on most of these chromosomes were characterized by high population differentiation and co-located with several GEA outliers strongly associated with environmental variables except for Freshwater Drum, where no GEA outliers were found within clusters of HMM islands. In all six chromosomes with clustered architectures we found, as expected, significantly higher Fstp values within the HMM islands (Figure 6). Comparisons of *H_O_* and *D_xy_* between HMM islands and chromosomal background showed three different patterns among six chromosomes: (1) islands on chromosome 7 and 17 in Freshwater Drum and chromosome 9 in Emerald Shiner had similar *H_O_* and *D_xy_*; (2) islands on chromosome 13 in Channel Catfish had significantly higher values of *H_O_* and *D_xy_*; (3) islands on chromosome 20 and 28 in Channel Catfish had significantly lower values of *H_O_* and *D_xy_* (Figure 6). We also found significantly elevated LD within the islands in all chromosomes except for chromosome 13 in Channel Catfish (Figure 6). Taken together, these results indicate that the HMM islands on the six chromosomes with clustered architectures have higher relative divergence than their chromosomal backgrounds; the islands on chromosome 13 in Channel Catfish also demonstrated higher absolute divergence, though without elevated LD.

**Figure 6.**
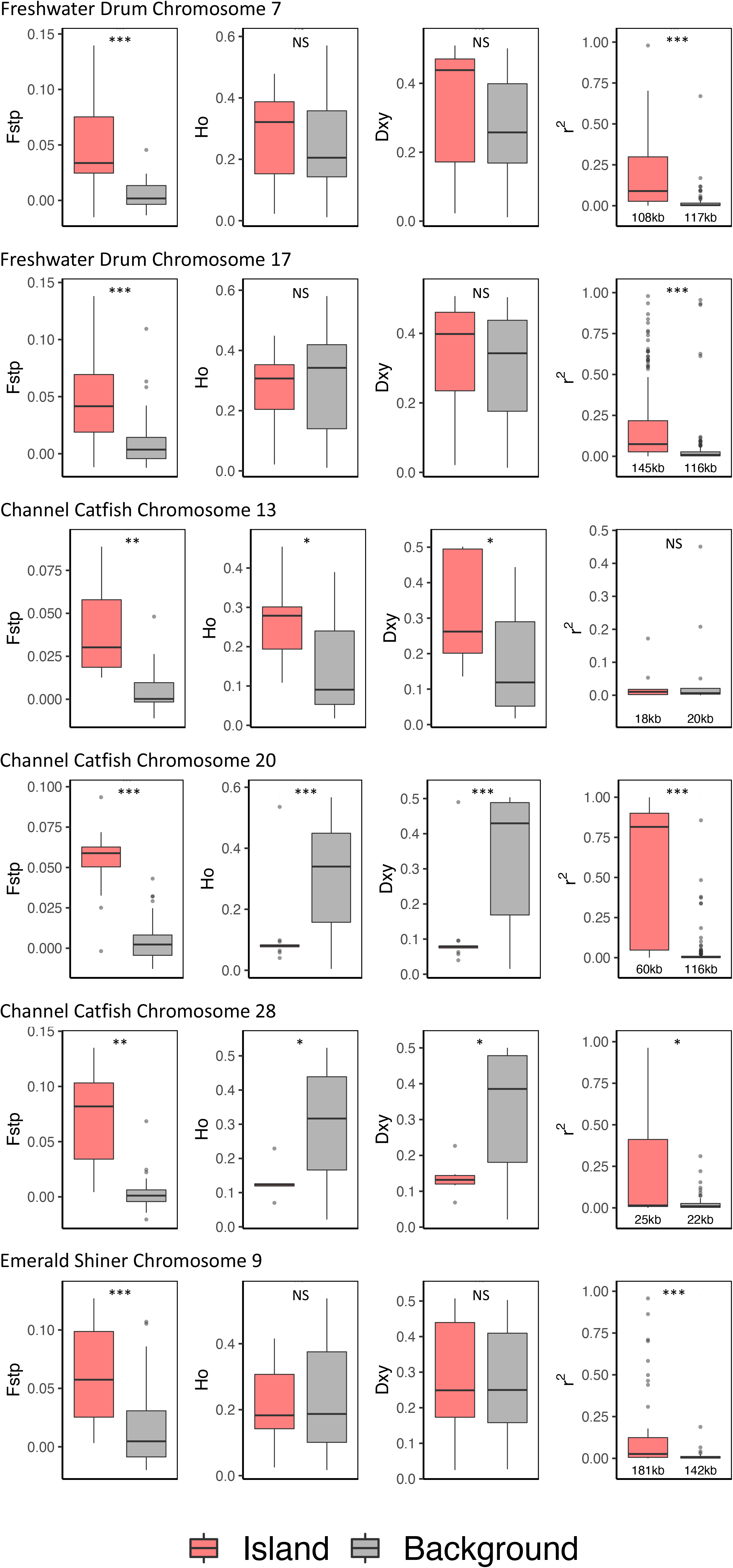
Comparisons of corrected *F*_ST_ (Fstp), heterozygosity (*H_O_*), absolute divergence (*D_xy_*), and LD (r^2^) between SNPs within the HMM islands (Island, red) and five combined random windows (Background, gray) on the corresponding chromosomes for chromosomes with clustered architecture, including on chromosome 7 and 17 in Freshwater Drum, chromosome 20 and 28 in Channel Catfish, and chromosome 9 in Emerald Shiner. The average distance between pairs of SNPs within islands and random windows were labelled below the bloxplots of LD values (column 4). Significance was assessed between islands and random windows using Wilcoxon tests with alpha level of 0.05. Note: *** = 0.001, ** = 0.01, * = 0.05, NS = not significant. Only HMM islands with more than one SNP were included.

A total of 9, 12, and 2 GO terms were significantly enriched (*p* < 0.05) in the HMM islands on chromosome 17 in Freshwater Drum, the island on chromosome 28 in Channel Catfish, and the inversion on chromosome 9 in Emerald Shiner, respectively (Table S7). Enriched GO terms included regulation of cellular component size, cell communication, and regulation of ion transmembrane transport. There were no annotated genes found within the HMM island(s) on chromosome 7 in Freshwater Drum, chromosome 13 and 20 in Channel Catfish.

## Discussion

### Neutral Population Structure Reflects Differences in Life History Strategies Among Species

We found highly variable neutral population structure among our six riverine fish species that generally reflected differences in life history strategies. For example, both Bullhead Minnow and Bluegill, which had the highest levels of genetic differentiation, are nest spawners whose eggs and larvae are not transported by currents, limiting gene flow. In contrast, Gizzard Shad and Emerald Shiner, which had the lowest levels of structure in our study, are both broadcast spawners, allowing their eggs to be carried freely by the currents, facilitating gene flow. Genetic studies on similar fish species have generally corroborated the patterns we observed, with nest spawning species such as smallmouth bass (*Micropterus dolomieu*) exhibiting high levels of genetic structure in open systems compared to broadcast spawning species such as walleye (*Sander vitreus*) (Ruzich *et al*. 2019; Euclide *et al*. 2020; 2021)

An exception to the pattern described above was Channel Catfish, as they are nest spawners but displayed relatively low levels of differentiation. It is possible that the highly migratory nature of this species mixed with potentially low spawning fidelity (Pellett *et al*. 1998) could explain the low to intermediate levels of population differentiation we observed. Freshwater Drum also deviated from the expected patterns of population structure based on life history, as they are migratory broadcast spawners but displayed an intermediate level of population structure, with individuals from La Grange along with some individuals from southern populations in Pool 26 and Open River forming a distinct group. One possible explanation for this pattern is limited movement of Freshwater Drum between the Illinois River, where La Grange is located, and the mainstem Mississippi River. Unfortunately, movement data for this species are generally lacking, making it difficult to corroborate this hypothesis without additional research.

### GEA Outliers Reflect Adaptive Divergence in Response to Habitat Heterogeneity

Most of the GEA outliers that we found were associated with environmental PC1, which had the highest loadings for temperature and turbidity. It is likely that these GEA outliers reflect adaptive divergence driven by the large latitudinal gradient that we sampled. Our study system spans two major Köppen climate zones, with pools 4, 8, and 13 in a humid continental climate characterized by warm summers and very cold winters (below 0 °C), and Pool 26, Open River, and La Grange in a humid subtropical climate characterized by very warm and humid summers and mild winters (above 0 °C). Although we could not disentangle the effects of temperature and turbidity because they co-varied, we suspect that temperature is likely a major selective force shaping adaptive divergence in our study system given its pervasive effects across all levels of biological processes, from the biochemistry of metabolism (Deutsch *et al*. 2015) to reproduction (Pankhurst & Munday 2011) and the fact that most fish are ectotherms. Multiple studies have illustrated strong signals of adaptive divergence across temperature gradients in continuously distributed marine species, even when differentiation at neutral markers is low (Limborg *et al*. 2012; Stanley *et al*. 2018; Wilder *et al*. 2020). However, few studies have investigated temperature-mediated adaptive divergence in continuously distributed freshwater fish. Our study suggests riverine fish display patterns of strong adaptive divergence driven by temperature that are similar to those found in marine systems, highlighting the fact that populations of continuously distributed riverine species may display the potential for local adaptation across their range.

While GEA outliers for most species in our study were generally associated with environmental PC1, outliers in Freshwater Drum were associated with environmental PC2, which displayed high loadings for measures of productivity including chlorophyll and nitrogen, and to a lesser extent, flow. This result suggests that the environmental variables influencing adaptive divergence in Freshwater Drum may differ from our other study species. Specifically, it is possible that Freshwater Drum is more affected by eutrophication caused by agricultural runoff compared to our other study species. Numerous studies have demonstrated that fish species respond differently to eutrophication depending on their life histories (Tammi *et al*. 1999; Hondorp *et al*. 2010; Jacobson *et al*. 2017). Alternatively, Freshwater Drum might have evolved in response to an underlying geomorphological condition correlated with agricultural inputs or to variation along a lotic-lentic gradient, to which Freshwater Drum are known to respond (Rypel *et al*. 2006; Jacquemin *et al*. 2015).

### Gene Flow Influences the Genomic Architecture of Local Adaptation

Theoretical studies and genetic simulations predict that increased gene flow will lead to increasingly concentrated genomic architecture of adaptation (Yeaman & Whitlock 2011; Via 2012; Yeaman 2013). However, few empirical studies have tested this hypothesis in natural populations, and the results of these empirical studies have not necessarily supported theoretical work (Burri *et al*. 2015; Renaut *et al*. 2019). Our study included six fish species spanning a wide gradient of genetic differentiation (overall *F*_ST_ from 0.0004 – 0.07), indicating highly variable levels of gene flow. Gene flow appeared to be correlated with the landscape of adaptive divergence, as species with high gene flow (Emerald Shiner, Channel Catfish and Freshwater Drum) displayed more clustered architecture of adaptation than low gene flow species (Bullhead Minnow and Bluegill). Our results are somewhat similar to a recent study which examined adaptive divergence of four flatfish species across a strong salinity gradient in the Baltic Sea (Le Moan *et al*. 2019). Specifically, Le Moan *et al*. (2019) found more evidence of clustered architectures of adaption in species displaying low genetic differentiation compared to those displaying higher differentiation. However, Le Moan *et al*. (2019) sampled a much smaller gradient of genetic differentiation (overall *F*_ST_ from 0.005 – 0.02) than our study and examining the effects of gene flow on landscapes of adaptive differentiation was not a central goal of their study.

Though our finding that clustered genomic architectures of adaptation (i.e., genomic islands of divergence) increase with gene flow is in line with theoretical expectations and the results from Le Moan *et al*. (2019), this finding is inconsistent with other studies positing that islands of divergence are the result of variation in intrinsic recombination rate rather than the combination of gene flow and selection (Roesti *et al*. 2012; Renaut *et al*. 2019). In fact, there is considerable debate over the mechanisms that lead to islands of divergence, with past research suggesting that these islands can be caused by variation in recombination rates (Roesti *et al*. 2012; Renaut *et al*. 2019), linked selection (Cruickshank & Hahn 2014; Burri *et al*. 2015), divergence hitchhiking (Via 2012), genomic rearrangements including chromosomal inversions (Rogers *et al*. 2013; Yeaman 2013), and elevated linkage preserving locally adapted alleles (Yeaman & Whitlock 2011). While the cluster of islands on chromosome 9 in Emerald Shiner appears to be caused by an inversion (see following section), the mechanisms that created the other islands are less clear.

To investigate the genomic mechanisms that created the islands on the remaining five chromosomes exhibiting clustered architecture, we calculated the following four metrics: LD, *F_ST_, H_O_*, and *D_xy_*, and compared these metrics between islands and chromosomal background. While islands on all five chromosomes displayed elevated *F*_ST_ as expected, we did observe differences in the remaining three metrics among the chromosomes. Islands on all but one chromosome displayed elevated LD; *H_O_* was elevated or similar to neighboring neutral regions in islands on three out of five chromosomes, and *D_xy_* was elevated or similar to neutral regions in islands on the same three chromosomes. While LD can be a useful metric for understanding genomic processes, we found that it did not help us differentiate the mechanisms responsible for creating islands in the current study and instead focused on *H_O_* and *D_xy_*. Estimates of *H_O_* and *D_xy_* suggest that the islands on the two chromosomes with reduced diversity (islands on Channel Catfish chromosomes 20 and 28) may have been created by linked selection (Cruickshank & Hahn 2014; Burri *et al*. 2015), while the islands on Channel Catfish chromosome 13 and Freshwater Drum chromosomes 7 and 17 may have arisen through divergent selection (Kulmuni & Westram 2017).

Islands created by divergent selection are hypothesized to have a major role in facilitating adaptive divergence with gene flow, whereas islands created by linked selection are likely a result of the underlying genomic landscape and do not necessarily reflect recent adaptive divergence (Cruickshank & Hahn 2014; Burri *et al*. 2015). Thus, it is extremely important to differentiate these two types of islands when investigating adaptive divergence. The most effective way distinguish between these island types is to compare measures of absolute diversity, as islands created by linked selection should show reduced absolute diversity while islands created by divergent selection should not (Cruickshank & Hahn 2014; Irwin *et al*. 2016). Applying this method to our data provided evidence that islands on three of the chromosomes in our study were created by divergent selection and are likely involved in adaptive divergence with gene flow whereas the islands on the other two chromosomes were likely a result of ancient linked selection that acted to reduce diversity in particular genomic regions but is not influencing contemporary adaptive divergence.

Interestingly, we observed variation in the mechanisms putatively responsible for creating islands both within and among species, as islands on one of the three chromosomes in Channel Catfish were likely created by divergent selection while islands on the other two chromosomes were likely the result of linked selection. This result indicates that, while many studies have tried to generalize the proposed mechanisms responsible for creating islands of divergence, these mechanisms vary across and even within species (Ottenburghs *et al*. 2020; Liu *et al*. 2020; Wilder *et al*. 2020). Thus, it is vital to examine both absolute and relative measures of differentiation as well as diversity in newly discovered islands of divergence to help clarify the mechanisms responsible for creating these islands.

### A Chromosomal Inversion Facilitates Local Adaptation with High Gene Flow in Emerald Shiner

Our results and those of previous empirical and theoretical studies suggest that divergent selection can result in clusters of adaptive loci through mechanisms such as divergence hitchhiking when gene flow is relatively high (Yeaman & Whitlock 2011; Via 2012). However, when gene flow is extremely high, it is likely that additional genomic mechanisms, such as structural polymorphisms, may be required to protect clusters of adaptive loci from among-population recombination caused by gene flow (Yeaman & Whitlock 2011; Rogers *et al*. 2013; Yeaman 2013; Tigano & Friesen 2016). The gradient of gene flow sampled in our study presents an excellent opportunity to test this hypothesis. In our study, clustered architectures of adaptation were common in species with relatively high gene flow, such as Channel Catfish and Freshwater Drum (average overall *F*_ST_ = 0.004), but these clustered architectures did not appear to be associated with structural polymorphisms. In contrast, in Emerald Shiner, the species with highest gene flow (overall *F*_ST_ = 0.0004), nearly all of the adaptive loci identified were found in a single genomic region that displayed strong evidence of a chromosomal inversion. Taken together, our results provide novel empirical evidence to support the theory that chromosomal inversions are important for facilitating adaptive divergence in systems with extremely high gene flow.

Our study also adds to the growing body of evidence that chromosomal inversions are important for facilitating adaptive divergence in continuously distributed fish species. Inversions putatively involved in adaptive divergence have been documented in many fishes including Atlantic cod (*Gadus morhua*) (Kirubakaran *et al*. 2016), lingcod (*Ophiodon elongatus*) (Longo *et al*. 2020), rainbow trout (*Oncorhynchus mykiss*) (Arostegui *et al*. 2019; Pearse *et al*. 2019), Pacific herring (*Clupea pallasii*) (Petrou *et al*. 2021), Atlantic silverside (*Menidia menidia*) (Wilder *et al*. 2020), and European plaice (*Pleuronectes platessa*) (Le Moan *et al*. 2019). However, all of these studies were conducted on marine fish or salmonids, making our study the first to provide evidence of a putative adaptive inversion in a non-salmon freshwater fish. It is likely that the lack of previous evidence for adaptive inversions in freshwater fish is due to the generally higher genetic structure observed in these species, making inversions less necessary for adaptation. However, our study illustrates that inversions are likely a larger component of adaptive divergence in freshwater fish than previously assumed, highlighting the importance of future studies aimed at characterizing them in additional species.

Although inferring the functional significance of the putatively adaptive inversion that we detected is difficult, it is possible to speculate on its role in facilitating adaptive divergence. The putatively derived variant of this inversion was only detected in the three southern river reaches in our study, which are substantially warmer and more turbid than northern reaches. This suggests that the derived inversion variant may have evolved and increased in frequency as Emerald Shiner adapted to warmer and/or more turbid environments in more southern regions. Inversions putatively linked to adaptive divergence across environmental and latitudinal gradients have also been identified in marine species such as lingcod (Longo *et al*. 2020) and Atlantic silverside (Wilder *et al*. 2020), but these studies faced similar difficulties when attempting to describe the functional significance of the adaptive inversions they identified. Future research combining whole genome resequencing with physiological challenge studies would be useful for assessing the functional role of these inversions in the process of adaptive divergence.

## Conclusions

Our data from six riverine fish species in UMRS displaying a large gradient of life history strategies suggest that higher gene flow leads to increasingly concentrated genomic architectures of adaptation. Additionally, our results provide evidence that the mechanisms that create islands of divergence can be highly variable across and within species, with both ancient linked selection and more contemporary divergent selection playing important roles in creating genomic islands of differentiation. Additionally, our study provides further evidence that chromosomal inversions are important for facilitating adaptive divergence in continuously distributed species with extremely high gene flow and also sheds light on the documented importance of inversions in freshwater fish. Taken together, our findings represent a significant contribution towards understanding the evolutionary processes that influence the genomic landscape of adaptation in non-model organisms. However, our study used RADseq, which does not assess the full suite of adaptive divergence across the genome. Future studies should focus on whole genome resequencing to better understand variation within genomic islands of divergence and to assess the functional role of these islands in promoting adaptive divergence.

## Acknowledgements

This study was funded by the U.S. Army Corps of Engineers Upper Mississippi River Restoration Program (grant number 96514790661277) and supported by the Turing High Performance Computing cluster at Old Dominion University. We thank U.S. Geological Survey Upper Midwest Environmental Sciences Center, Iowa Department of Natural Resources, Minnesota Department of Natural Resources, Wisconsin Department of Natural Resources, Illinois Natural History Survey, and Missouri Department of Conservation for their great efforts in sample collection. We thank Kristen Gruenthal for assistance in the laboratory, and Kathi Jo Jankoswki for help with environmental data access. We are grateful to Uland Thomas for allowing us to use his fish profile photos of the six study species and Hyeon Jeong Kim for help with photo editing. Any use of trade, firm, or product names is for descriptive purposes only and does not imply endorsement by the U.S. Government.

## Data Accessibility

Upon acceptance, demultiplexed RAD sequencing data used in this study will be released in the NCBI with BioProject ID, PRJNA674918. Bioinformatic scripts along intermediate datasets (post-filtering *vcf*, neutral *vcf*, adaptive *vcf*) supporting this article will be available as a Github repository.

## Author Contributions

YS, KLB, AB and WAL conceived of the study, designed the study, and coordinated the study. YS and WD carried out the molecular lab work. YS and WAL conducted data analyses and drafted the manuscript; GJM helped interpreted the results regarding putative inversions. MC supervised the project. All authors commented the manuscript and gave final approval for publication.

