## Supplementary Methods for "Gene flow influences the genomic architecture of local adaptation in six riverine fish species"

Yue Shi^*^, Kristen L. Bouska, Garrett J. McKinney, William Dokai, Andrew Bartels, Megan V. McPhee, Wesley A. Larson

**This PDF file includes:**

Supplementary Methods;

Supplementary References;

Figure S1 - S5;

**Other Supplementary Materials for this manuscript include the following:**

Table S1 - S7 (Excel);

**Supplementary Methods**

***RAD Sequencing and SNP Filtering***

DNA was isolated from fin clip samples preserved in > 95% ethanol with Qiagen DNeasy Blood and Tissue Kits. DNA extracts were quantified and normalized to 200 ng in 10 ul. Restriction site-associated DNA (RAD) libraries were prepared following the BestRAD protocol (Ali *et al.* 2016) using restriction enzyme *SbfI* and NEBNext^®^ Ultra^TM^ DNA Library Prep Kit for Illumina^®^ as detailed in (Ackiss *et al.* 2020). All prepared libraries were sent to Novogene (Sacramento, CA) for sequencing on the Illumina NovaseqS4 platform (PE 150).

Raw RAD sequences were processed using Stacks v2.3 (Rochette *et al.* 2019) following the protocol detailed in (Ackiss *et al.* 2020) with the following modifications: 1) PCR clones were removed using *clone_filter* prior to *ustacks*; 2) we changed the -M parameter in *ustacks* from 5 to 3 because our species do not have duplicated genomes. Initial SNP filtering was performed with VCFtools v0.1.16 (Danecek *et al.* 2011) and included 1) removing loci genotyped in fewer than 70% of individuals, 2) removing individuals missing more than 50% of loci, and 3) removing loci with a minor allele count less than 3. In addition, we used HDPlot (McKinney *et al.* 2016) to remove any putatively paralogous loci, which were loci with heterozygosity greater than 0.55 or a read ratio deviation greater than 5 and less than -5. Only the SNP with the highest minor allele frequency on each RAD tag was included in the final dataset. Putative sample duplicates were identified by calculating relatedness between samples using VCFtools (--relatedness). Relatedness was estimated only using loci with minor allele frequency (MAF) larger than 0.05 to avoid upward bias and putative sample duplicates were removed if relatedness levels were larger than 0.90. Finally, we assessed heterozygosity in individual samples to check for potential sample contamination. File format conversion was performed using PGDSpider v2.1.1.5 (Lischer & Excoffier 2011).

***Identification of Putatively Neutral and Adaptive Loci***

We used four differentiation-based outlier methods and three genotype-environment association (GEA) methods to identify (1) a dataset of putatively neutral SNPs and (2) putatively adaptive loci associated with environmental variables for each species. The differentiation-based outlier methods we used were Bayescan v2.0 with prior odds of 1000 (Foll & Gaggiotti 2008), Arlequin v3.5.2.2 (Excoffier & Lischer 2010), OutFLANK v0.2 (Whitlock & Lotterhos 2015) and R package *pcadapt* v.4.3.1(Luu *et al.* 2016). To initialize the hierarchical island model in Arlequin, we set the three upper reaches Pools 4, 8, 13 as one group, and the three lower reaches Pool 26, Open River and La Grange as another based on the most common pattern of population structure in the dataset (Fig. 1C). To run *pcadapt*, we determined K values using scree plot. The GEA methods we used were redundancy analysis (RDA) using the R package *vegan* v2.5-6, latent factor mixed model (LFMM2) in the R package *lfmm* (Caye *et al.* 2019) and Bayenv2 (Günther 2013). To run RDA, we used a three standard deviation cutoff to list loci with extreme loading scores on the significant RDA axes (1000 permutations, *p* < 0.05) as suggested by (Forester *et al.* 2018). To apply LFMM2, we first performed PCA and used broken stick model to determine the number of latent factors or K, then applied LFMM algorithm (lfmm_ridge and lfmm_test), and calibrated p values using genomic inflation factor (GIF). To run Bayenv2, we first generated covariance matrix of relatedness using all SNPs by taking the average of final matrices from 10 MCMC runs using the default run parameters. Bayes factor (BF) and Spearman rank correlation coefficient values (ρ) were calculated by taking the median of 10 independent runs of Bayenv2. Loci with top 1% BF values (BF > 3) and top 5% of ρ were considered as outliers. We adjusted the resulting *p* values for the false discovery rate (FDR) for all analyses when applicable by computing *q* values with the R package *qvalue* v2.16.0 (Lai 2017). A locus was considered as an outlier if *q* < 0.05.

**Table** Parameter settings in PCAdapt and LFMM in this study

| **Species** | **K (PCAdapt)** | **K (LFMM)** | **Adjusted GIF (LFMM)** |
| --- | --- | --- | --- |
| Bullhead minnow | 3 | 3 | 3 |
| Bluegill | 2 | 1 | 4.2 |
| Freshwater drum | 1 | 1 | 1.1 |
| Channel catfish | 2 | 1 | 1.2 |
| Gizzard shad | 1 | 1 | 1.2 |
| Emerald shiner | 2 | 1 | 1 |

***Functional Enrichment***

We conducted Gene Ontology (GO) enrichment tests to test for functional enrichment of genes in the HMM islands located within the six chromosomes displaying clustered architecture, including (1) chromosome 7 and 17 in freshwater drum. (2) chromosome 13, 20, and 28 in channel catfish; (3) chromosome 9 in emerald shiner. We downloaded gene feature tables and GO information (version: Ensembl Genes 101) of the corresponding mapping reference genomes (Table S5) using NCBI and *biomaRt* v2.40.5 package in R (Durinck *et al.* 2005; 2009). We extracted genes within 10 Kb of a SNP for all SNPs located within the islands for all six chromosomes except for chromosome 9 in emerald shiner. Since all of the HMM islands on chromosome 9 in emerald shiner were clustered inside of the identified inversion and there was relatively a smaller number of aligned loci, we extracted genes within 20 Kb of a SNP for all SNPs located within the identified inversion on the chromosome 9 in emerald shiner instead. See Supplementary Materials for detailed methods about GO enrichment tests. We tested functional enrichment of GO terms (Biological Process category only) in the regions of interest using *topGO* v2.36.0 package in R (Alexa *et al.* 2006) with a node size of 5 and the default “weight01” algorithm to account for the hierarchical structure of GO terms. We used α = 0.05 to determine significance and reported *p* values without multiple testing correction as suggested by the authors of *topGO*.

**Supplementary Figures**

**Figure S1.** Principal component analyses using 20 environmental variables for the six study reaches. A) The first two PCs are retained based on Kaiser-Guttmann criterion and broken stick model. Contribution of each environmental variable to PC1 and PC2 are shown in (B) and (C).

**Figure S2.** LD decay as a function of the distance (Kbp) separating two loci for chromosomes with outlier regions identified by R package *lostruct* with discreteness > 0.89 and where the middle PCA cluster showed significantly higher heterozygosity than the other two clusters. (A) chromosome 2, 8, and 21 in bullhead minnow; (B) chromosome 1, 8, and 12 in freshwater drum; (C) chromosome 2, 10, 12, 13, 14, and 16 in channel catfish; ; (D) chromosome 1, 4, 6, 10, and 13 in gizzard shad; (E) chromosome 6, 9, and 19 in emerald shiner.

**Figure S3.** LD heatmaps for chromosome 14 in Channel Catfish using all individuals (left) and only individuals homozygous for the more common orientation (right).

**Figure S4.** Characterization of putative inversion on chromosome 6 in Emerald Shiner. (A) PCA based on SNPs within the putative inversion region. Three clusters identified using k-means clustering correspond to two homozygote groups (blue and red) and a heterozygote group (purple). The discreteness of the clustering was calculated by the proportion of the between-cluster sum of squares over the total using the R function *kmeans* in *adegenet*. (B) Observed individual heterozygosity in each PCA cluster. Significance was assessed using Wilcoxon tests (α = 0.05). Note: *** = 0.001. (C) Genotype frequency distribution for the putative inversion across six study reaches. Bars represent the proportion of individuals belonging to a PCA cluster. (D) and (E) are LD heatmaps for chromosome 6 using all individuals (D) and only individuals homozygous for the more common orientation (E).

**Figure S5.** Characterization of putative inversion on chromosome 19 in Emerald Shiner. (A) PCA based on SNPs within the putative inversion region. Three clusters identified using k-means clustering correspond to two homozygote groups (blue and red) and a heterozygote group (purple). The discreteness of the clustering was calculated by the proportion of the between-cluster sum of squares over the total using the R function *kmeans* in *adegenet*. (B) Observed individual heterozygosity in each PCA cluster. Significance was assessed using Wilcoxon tests (α = 0.05). Note: *** = 0.001. (C) Genotype frequency distribution for the putative inversion across six study reaches. Bars represent the proportion of individuals belonging to a PCA cluster. (D) and (E) are LD heatmaps for chromosome 19 using all individuals (D) and only individuals homozygous for the more common orientation (E).
