## Supplementary figures and images for "Gene flow influences the genomic architecture of local adaptation in six riverine fish species"

### Figure S1

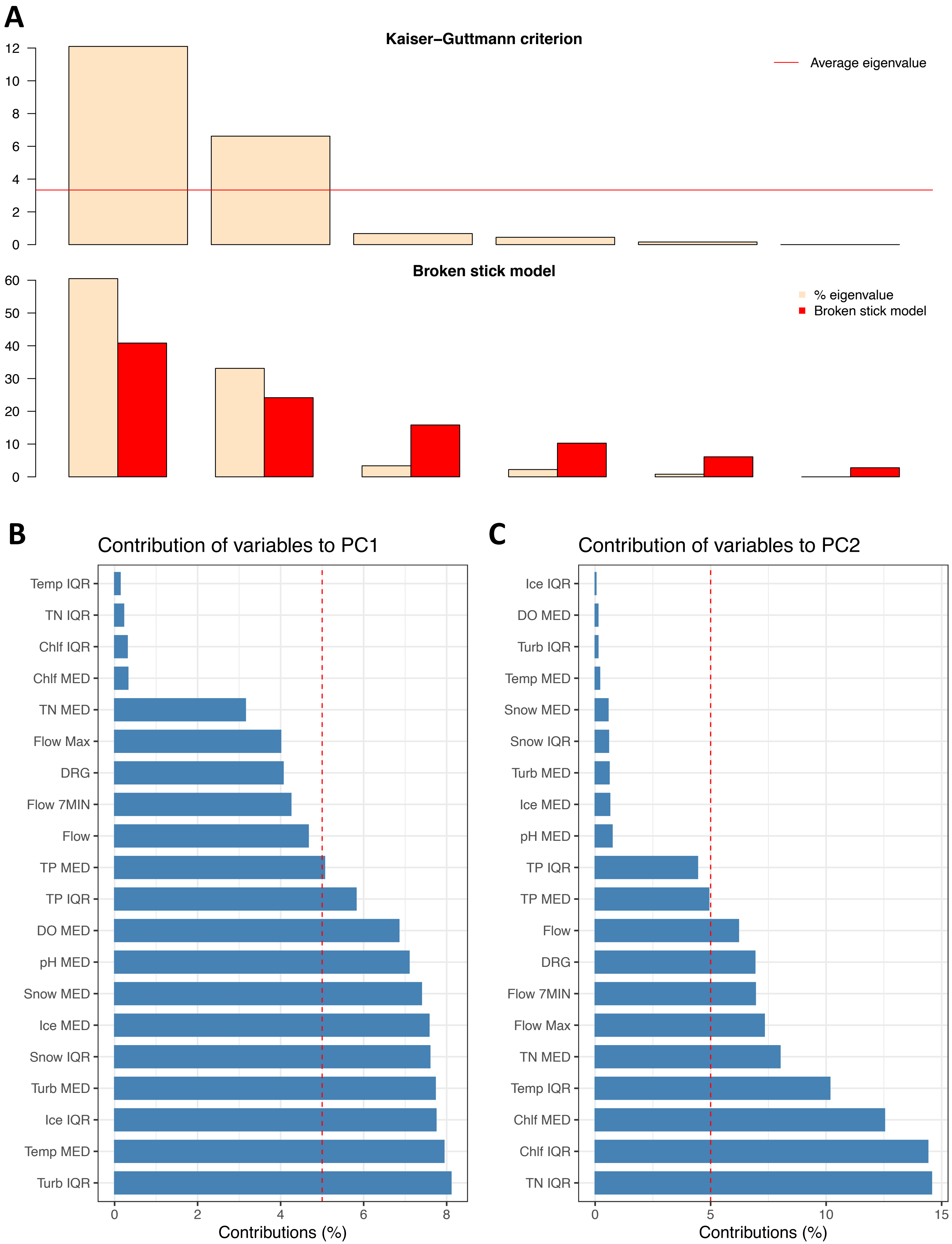

### Figure S2

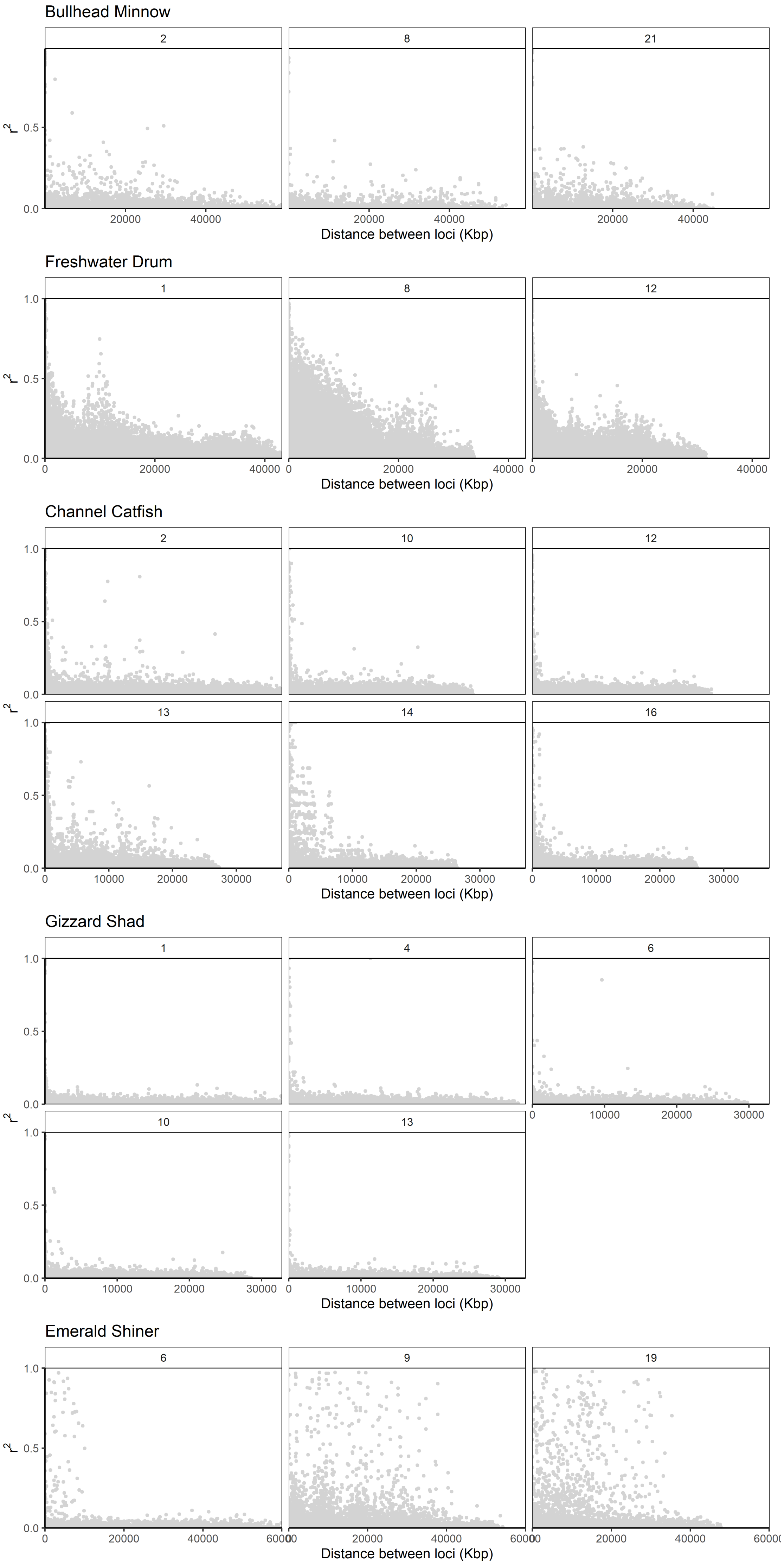

### Figure S3

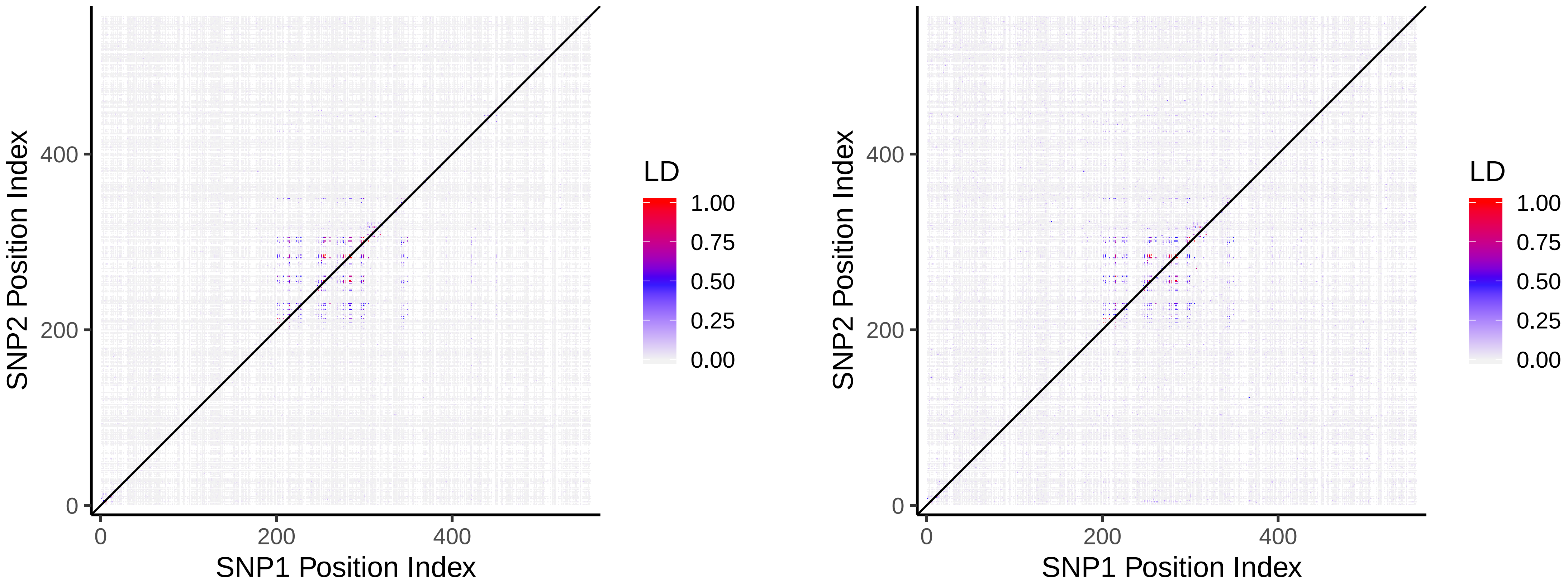

### Figure S4

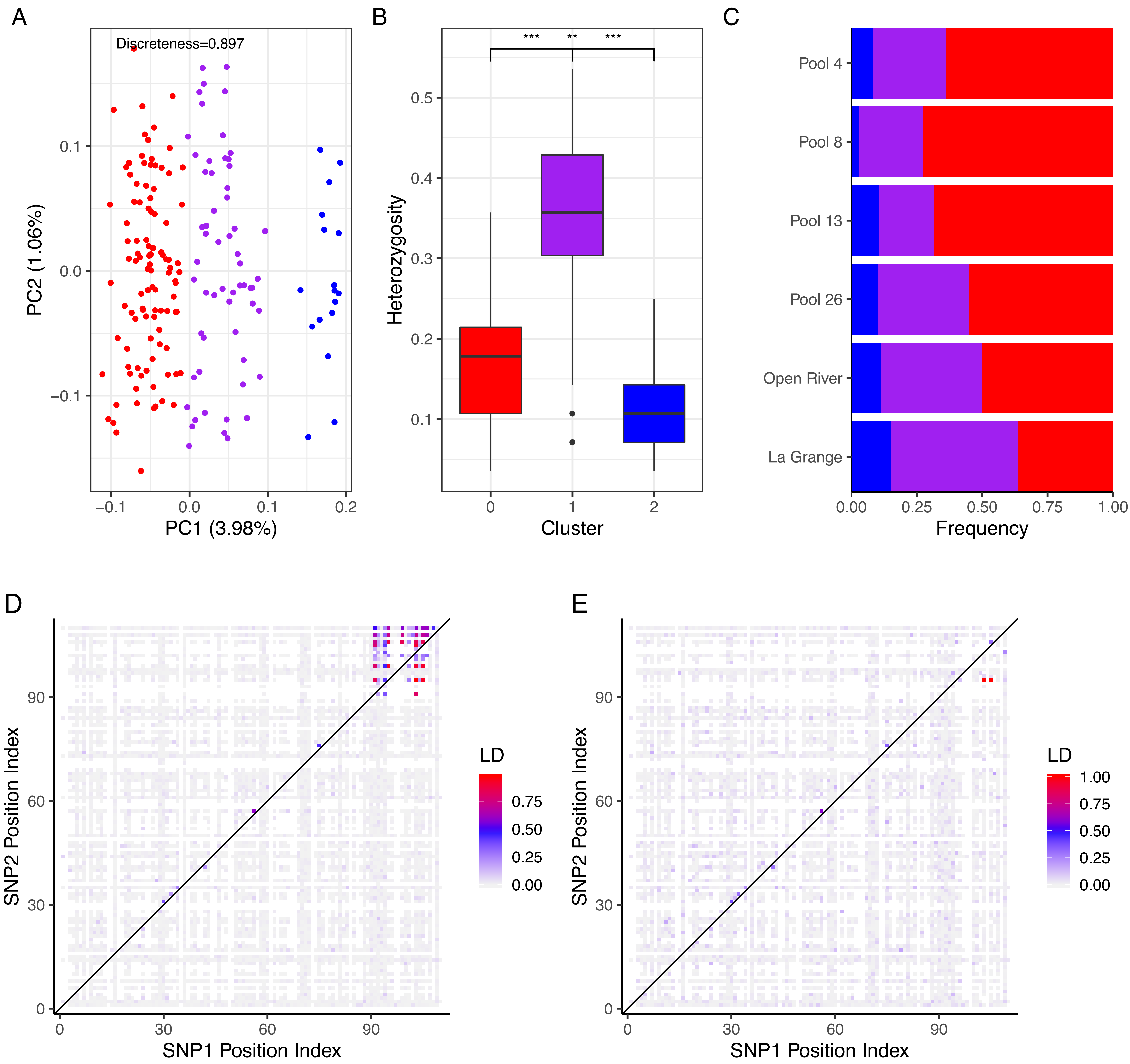

### Figure S5

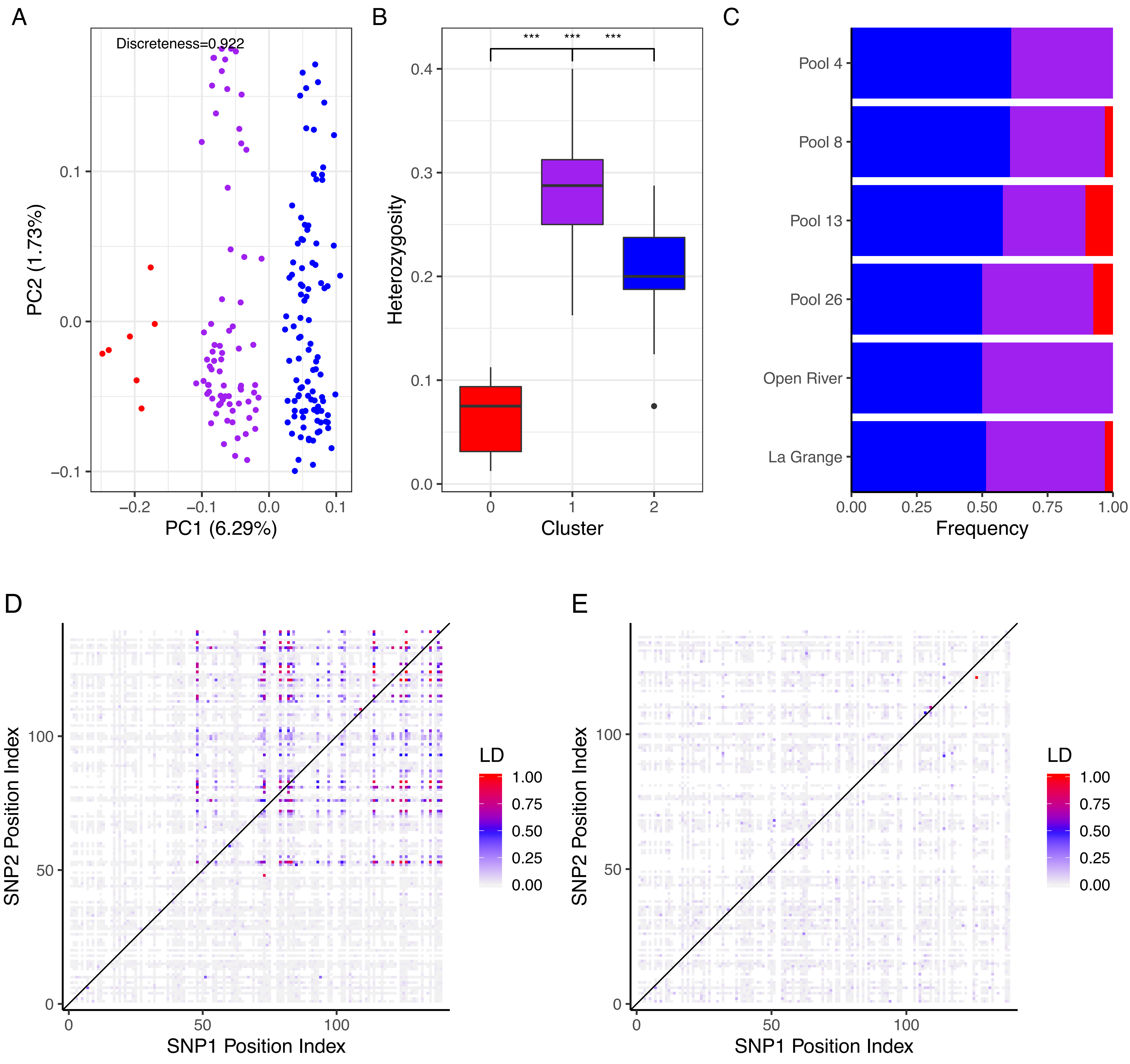
